# ReCo: automated NGS *re*ad-*co*unting of single and combinatorial CRISPR gRNAs

**DOI:** 10.1101/2023.03.09.530923

**Authors:** Martin Wegner, Manuel Kaulich

## Abstract

**Summary:** CRISPR screens are increasingly performed to associate genotypes with genotypes. So far, however, their analysis required specialized computational knowledge to transform high-throughput next-generation sequencing (NGS) data into sequence formats amenable for downstream analysis. We developed ReCo, a stand-alone and user-friendly analytics tool for generating read-count tables of single and combinatorial CRISPR library and screen-based NGS data. Together with cutadapt and bowtie2 for rapid sequence trimming and alignment, ReCo enables the automated generation of read count tables from staggered NGS reads for the downstream identification of gRNA-induced phenotypes.

**Availability and implementation:** ReCo is published under the MIT license and available at: https://github.com/KaulichLab/ReCo.

**Supplementary information:** Supplementary data are available at Bioinformatics online.

## 1 Introduction

The CRISPR-Cas system has emerged as an important tool for genome editing (Wang and Doudna, 2023). In its engineered version, the system consists of two components, a Cas endonuclease and a single gRNA (sgRNA) that guides the Cas enzyme to a predefined locus in the genome. Depending on the type of system, the targeted locus can be perturbed in multiple ways, among them the induction of double-strand breaks (causing insertions or deletions, InDels), editing of individual bases (base or prime editing), or recruiting effector domains to activate or repress gene transcription (CRISPRi or CRISPRa) (Liu *et al.*, 2022). In its most widely used form, a Cas nuclease, e.g. SpCas9, induces a DNA double-strand break in coding exons resulting in frameshift mutations that cause functional knockouts of the genes of interest. When target sites are bundled in a gRNA library, a population of mutant cells can be generated and screened for a phenotype of interest, enabling unbiased genotype-to-phenotype associations (Bock *et al.*, 2022). To do so, the gRNA expression cassette is stably integrated into the host cell genome which allows its population frequency to be used as a surrogate for the gRNA-induced phenotype (Ford *et al.*, 2019). gRNA frequencies are quantified by NGS, comparing different screening time points with their gRNA library frequency. Due to the low sequence diversity of gRNA libraries (only the gRNA part of the NGS-read is variable), gRNA amplicons are commonly sequenced with staggered oligos, rendering the gRNA position random within a window of up to 8 nucleotides, which avoids low diversity issues during NGS runs. This, however, prevents the extraction of gRNA sequences from NGS reads in which the gRNA position is fixed which requires additional read trimming and alignment steps for data processing. Although this setup is widely used, there is a lack of automatic pipelines to generate gRNA read count tables from staggered NGS data that enable computationally less developed groups to analyze their CRISPR libraries and screening samples. Closing this gap, we present the <u>*Re*</u>ad <u>*Co*</u>unting tool ReCo that automatically generates read count tables from single-end and paired-end NGS fastq files with minimal input requirements.

ReCo is implemented as a Python 3 package that can also be run as a standalone command line tool. It uses the parallelization capabilities of two external tools, cutadapt and bowtie2 (Martin, 2011; Langmead and Salzberg, 2012), to decrease sample processing time. ReCo can process arbitrary numbers of single-end and paired-end samples per run, corresponding to single or combinatorial CRISPR gRNA libraries. The tool requires minimal information per sample, but a unique sample name, as well as fastq and gRNA library file locations. Optionally, ReCo integrates expected sequencing depths and accepts vector maps in SnapGene format to account for 3Cs-technology-based samples (Diehl *et al.*, 2021; Wegner *et al.*, 2019, 2020). If provided with a vector file, ReCo will automatically find the 3Cs-template sequences and report their abundance in the final report.

## 2 Results

PinAPL-py is the only open-access tool for generating read count files from staggered NGS runs (**Fig. 1A**). However, the implementation of PinAPL-py leaves room for improvement, particularly for samples with high sequencing depths or total reads (Spahn *et al.*, 2017). Additionally, PinAPL-py is unable to handle combinatorial CRISPR samples. Thus, we implemented the stand-alone command line tool ReCo to automatically trim, align, and count single and combinatorial gRNA sequences from staggered NGS reads. Unlike previous tools, ReCo demands little user interaction and runs locally without the need to upload data, for which the connection speed can be a limiting factor. ReCo operates based on provided sample names, Illumina fastq and gRNA library file locations. Users can optionally provide expected sequencing depths per sample as well as plasmid maps in SnapGene (.dna) format to identify and account for 3Cs-technology-derived samples. ReCo then identifies the 3Cs placeholder gRNA sequence by sample subsampling (Supplementary Figure S1). Sequence trimming and alignment by cutadapt and bowtie2, respectively, are implemented iteratively while providing the option to use parallelization parameters of both tools to decrease running time per sample. To improve run time further, particularly for highly diverse samples, all unique putative sequences are first counted and then aligned to the gRNA sequence library. As physical output, ReCo provides .csv read count files per sample containing the gRNA counts. For trouble-shooting purposes, ReCo reports all sequences that could not be aligned to the provided gRNA library. The final read-count table is visualized with a plot panel in .png and .pdf formats, and a .txt file reporting the ReCo run parameters. The analysis plot panel summarizes trimming and alignment rates, expected and observed sample sequencing depth, the distribution of read counts, as well as gRNA completeness and sample distribution skew. Optionally, the 3Cs template gRNA placeholder sequence is highlighted for quality control purposes.

**Figure 1:**
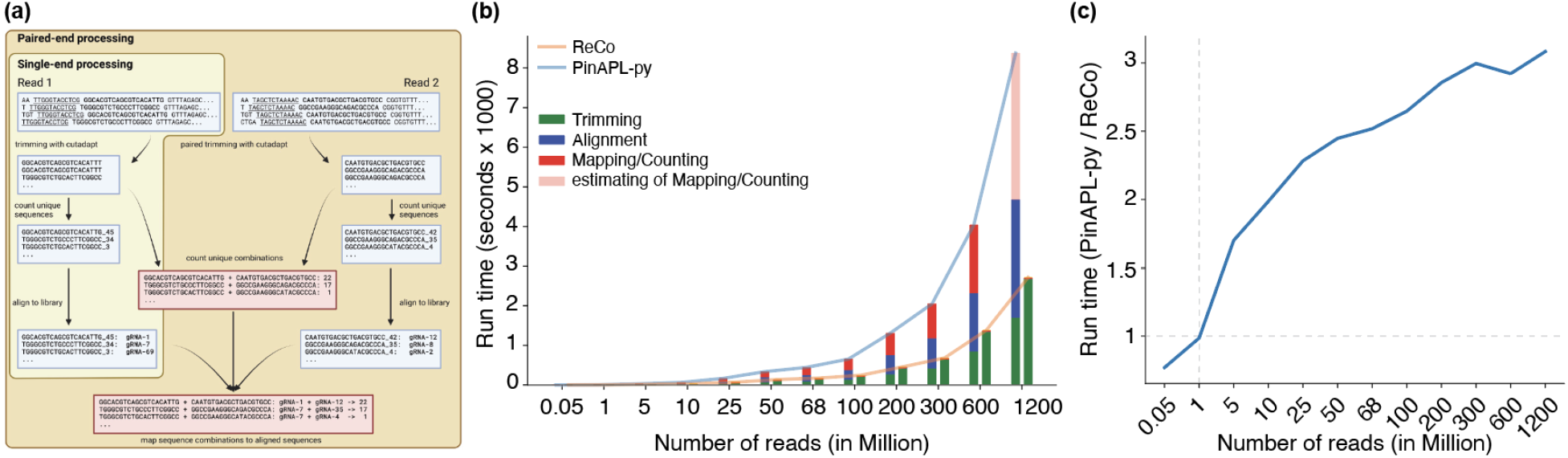
**(a)** Trimming and alignment strategy of ReCo. Single-end read samples are trimmed with cutadapt to isolate the putative gRNA sequences. Unique sequences are counted and then aligned only once to the gRNA library to reduce the number of required alignments. Paired-end read samples use the single-end pipeline for both reads individually. In an intermediate step, unique sequence combinations from corresponding reads are counted. In the final step, the unique gRNA combinations are mapped against the individual sets of alignments. The graph was created with BioRender.com. **(b)** Benchmarking of running times for ReCo and PinAPL-py. Sequencing samples of sizes between 0.5 million and 1.2 billion reads were processed and the time that was required for the individual steps was measured in seconds and is shown for each sample. While in the PinAPL-py algorithm, the time requirements grew for each processing step, in the ReCo algorithm, only the trimming procedure required more time in relation to the number of input reads. **(c)** The ratio of ReCo and PinAPL-py running times increases with the number of processed reads, meaning that the time requirements for PinAPL-py increase faster than those for ReCo.

## 3 Benchmarking

To assess the relative performance of ReCo, benchmarking against PinAPL-py was performed (Spahn *et al.*, 2017). PinAPL-py was chosen as it is the only other available tool to operate on staggered NGS reads. Moreover, benchmarking was limited to single-end NGS reads, as PinAPL-py does not accept paired-end NGS reads. To assess their relative performance, we used the test data set provided by PinAPL-py, that are derived from a genome-wide CRISPR-Cas knockout screen using the SpCas9 Brunello gRNA library in a drop-out screen in A375 melanoma cells, containing 67.9 million reads. We separated the benchmarking in two aspects, the number of found gRNAs and their associated alignment rates, and the required run time. Alignment rates and the number of found gRNAs were similar with 83.88% and 83.91%, and 98.58% (76,341 of 77,441) and 98.62% (76,372 of 77,441) for PinAPL-py and RecCo, respectively. However, we found an issue within PinAPL-py’s alignment parameters which resulted in the failure to detect 31 gRNA sequences that are the reverse complement of other gRNAs, an issue that does not occur with ReCo. To benchmark the required run time, sampled data sets corresponding to 500K, 1M, 5M, 10M, 25M, 50M, 100M, 200M, 300M, 600M and 1.2B reads were derived from the original test data and processed individually with no other jobs runnning by PinAPL-py and ReCo with the same number of cores to maximize parallelization. While the run time of PinAPL-py grew exponentially with sample size and was dominated by trimming, alignment, and mapping/counting, the run time of ReCo was determined solely by trimming, with alignment and mapping/counting being decoupled from sample size (**Fig. 1B**). With increasing sample size, the ratio between the required run time for PinAPL-py and ReCo increased (**Fig. 1C**), demonstrating that ReCo scales better with samples of high diversity, such as combinatorial libraries or multiple diverse samples.

## 4 Conclusion

ReCo is a scalable read-counting tool for single and combinatorial CRISPR gRNA library data. It automatically recognizes gRNA positions in staggered single and paired-end NGS reads, generates read count files for further data analysis, and provides a visual quality control report summarizing the percentage of aligned and trimmed reads, expected and obtained sequencing depth, as well as gRNA and sample distribution skew. Combined with downstream CRISPR analysis tools, experienced and inexperienced users can efficiently analyze the effects of gRNAs/gene phenotypes across diverse CRISPR screen conditions.

## Data availability

The data underlying this article are available in the article and in its online supplementary material.

---

**Supplementary Figure S1:**
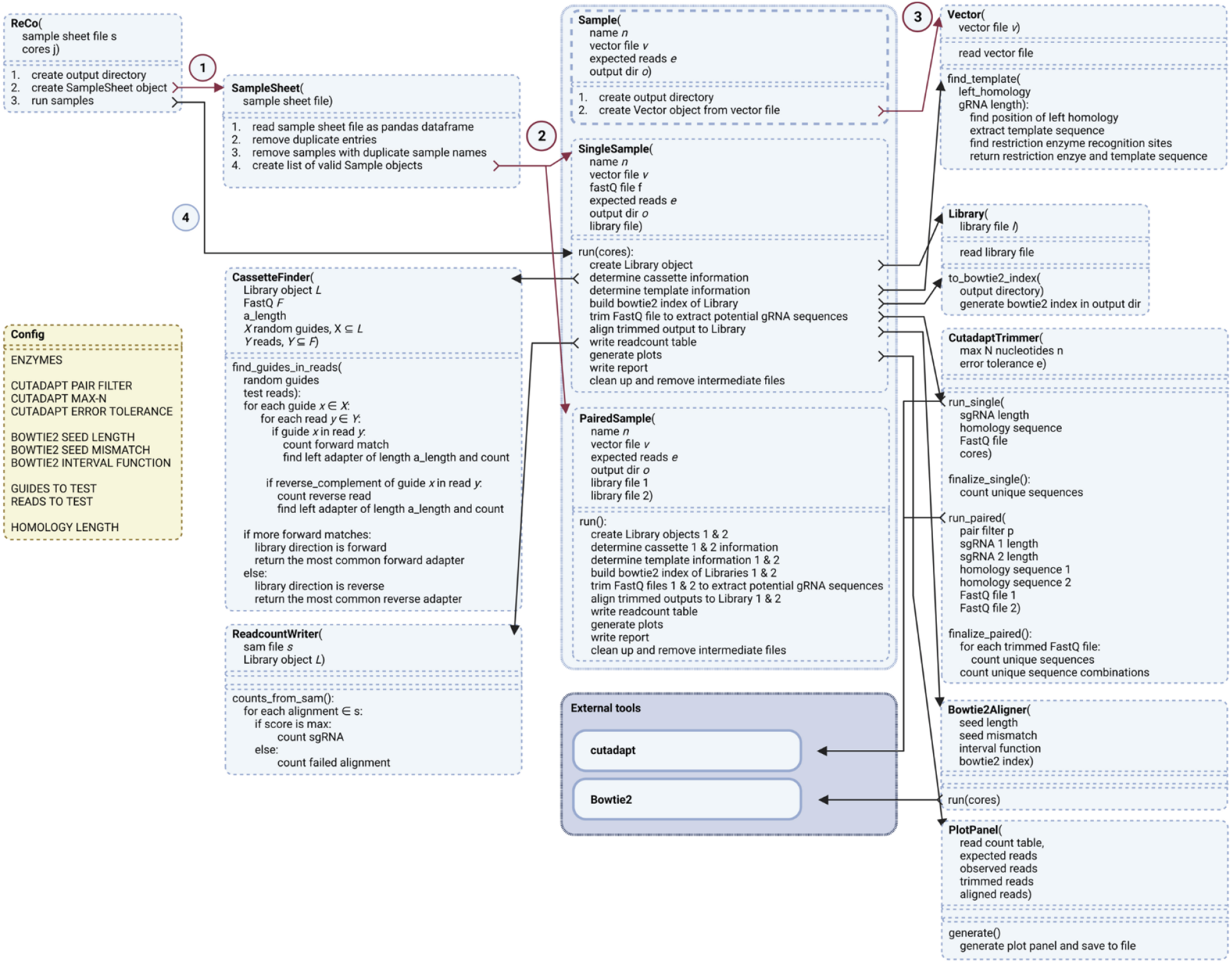
ReCo’s functional components are organized in a set of classes. The arrows and circled numbers 1 to 3 represent the initialization process. The black arrows and the circled number 4 represent the logical flow of processing a Single-end read sample (SingleSample). A Paired-end read sample (PairedSample) is processed analogously, for clarity the arrows are omitted. The Config class provides the configuration for ReCo and can be adjusted, the external tools cutadapt and bowtie are required to be available from the system path. The graph was created with BioRender.com.

**Supplementary Figure S2:**
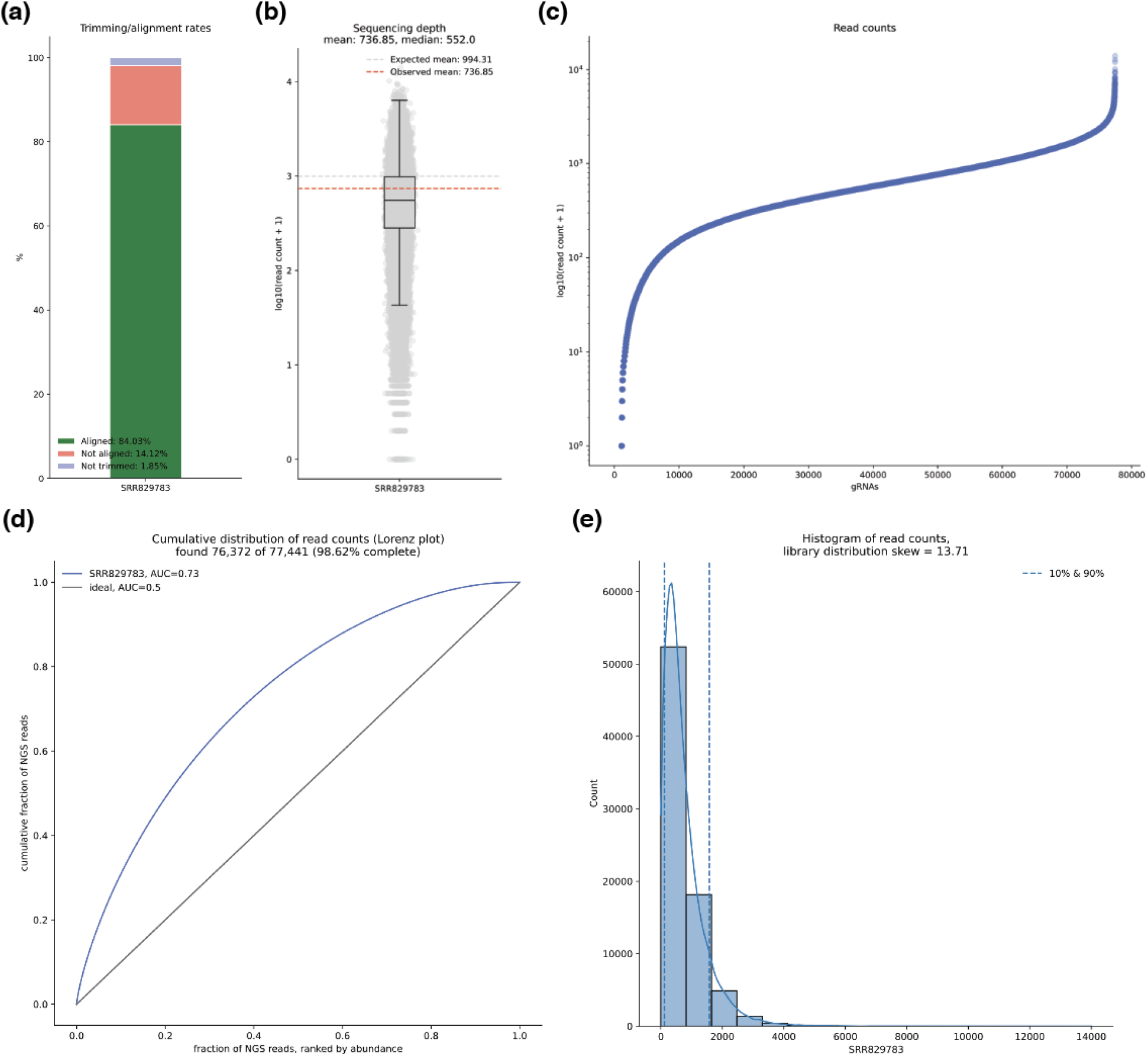
The ReCo plot panel visualizes the gRNA distribution of single-end and paired-end read samples. Here, a plasmid DNA preparation of the Brunello library (Doench *et al.*, 2016) was sequenced according to Spahn et al., 2017. The title of the plot contains the sample name, the number of expected reads, and the number of observed reads. **(a)** A stacked bar chart indicates the ratio of aligned (green color), not aligned (red color), and not trimmed (light blue color) reads. **(b)** A box plot displays the distribution of read counts on a logarithmic scale, the expected and observed mean read counts are highlighted with horizontal dashed lines in gray and red color, respectively. **(c)** The log-transformed distribution of read counts per gRNA, sorted increasingly. **(d)** A cumulative plot of gRNA abundance. The example library has an area under the curve (AUC) of 0.73. Out of a total of 77,441 gRNA sequences, 76,372 sequences were found, indicating that the library contains 98.62% of all intended gRNAs. For comparison, the cumulative distribution of an ideal, uniformly distributed library is plotted as a black diagonal line and has an AUC of 0.5. **(e)** A histogram of gRNA abundance and overlay of the corresponding density plot shows the skew of the library distribution. The 10 and 90 percentiles are indicated with vertical dashed lines, and the distribution skew of the library is 13.71.

